# Multi-trait colocalisation using *MystraColoc*: improved performance, deeper insights

**DOI:** 10.64898/2026.03.30.715409

**Authors:** Valentina Iotchkova, Michael E. Weale, Genomics Ltd

## Abstract

Multi-trait colocalisation is a vital tool to make sense of the large amounts of GWAS data available on platforms like Mystra. It identifies genetic association signals that cluster together, allowing us to infer which gene might be causal for a trait and also which constellation of biological effects might be affected by modulating that gene. Multi-trait colocalisation is a challenging computational problem. Here, we introduce *MystraColoc*, a Bayesian algorithm for multi-trait colocalisation that works across hundreds or even thousands of GWAS datasets. We illustrate its power both via a worked example at the *HDAC9*-*TWIST1* locus, and via a simulation study that demonstrates its superior clustering performance compared to alternative methods.

## Introduction

We are in the midst of a revolution in our use of data from genome-wide association studies (GWAS). Platforms like Mystra (https://www.mystra.com) provide access (in a harmonised and cross-referenced way) to tens of thousands of GWAS datasets. These collectively describe the statistical associations of millions of genetic variants in the human genome with thousands of traits or phenotypes, obtained from tens of thousands to millions of individuals sampled from around the world. The traits now available in such GWAS platforms range from molecular traits such as RNA (eQTLs) and protein (pQTLs) quantitative trait loci in different tissues and cell types, to high-level traits such as an individual’s susceptibility to disease or response to a drug.

A foundational type of analysis needed to make maximal use of such platforms is *colocalisation* - the identification of shared genetic signals between two or more studies representing different traits [1,2]. The identification of such shared genetic signals allows for the inference of shared genetic control, and therefore of shared biology. For example, the identification of a unique colocalisation between a genetic signal for disease A and gene expression for gene B in tissue C provides a hypothesis both about which gene is responsible for affecting disease A, and also about which tissue is important for this effect, both of which can aid in the development of novel drug targets. Likewise, the identification of a colocalisation between disease X and disease Y may either identify opportunities for drug indication expansion (if the putative risk allele increases the risk of both), or concerns for drug safety (if the putative risk allele increases the risk of one and decreases the risk of the other).

Methodologies for the colocalisation of two traits are relatively mature, and are represented by established methods such as *coloc* [1]. To extend colocalisation to multiple traits, one approach could be to apply a method like *coloc* to all trait pairs (perhaps restricting to only those traits with a significant genetic signal at the locus, to limit computational burden), and then apply statistical clustering techniques to the results. But this is far from ideal. Adding statistical clustering as a second stage does not make full use of the data, and traits with weak signals will be overlooked because the power from combining multiple datasets is lost.

True multi-trait colocalisation strategies have to contend simultaneously with more GWAS data and the quadratic combinatorial expansion of causality hypotheses among multiple studies. As a result, earlier methods for multi-trait colocalisation such as *moloc* [3] were in practice limited to no more than four traits. More recently, *HyPrColoc* [4] has emerged as a widely-adopted method, thanks to its ability to consider many more traits simultaneously. *HyPrColoc* is computationally efficient, partly because it approximates relevant posterior probabilities, and partly because it uses an irreversible ‘branch and bound’ clustering step to split traits into smaller clusters that are likely to share distinct causal variants.

Genomics has developed an alternative multi-trait colocalisation algorithm, *MystraColoc*, which adopts a different approach. Instead of using branch and bound divisive clustering, *MystraColoc* allows all cluster combinations to be considered through an efficient iterative Bayesian search method. Here, we illustrate the power of *MystraColoc*, first by considering a real-world example and then by a head-to-head comparison of *MystraColoc* and *HyPrColoc* under simulated conditions.

## Results

### *MystraColoc* applied to the *HDAC9-TWIST1* locus

There is a well known genetic association signal for cardiovascular disease and related traits that centres on the genetic variant rs2107595 [5]. This variant lies on chromosome 7, around 7kb downstream of *HDAC9* and 11kb upstream of *TWIST1* (Figure 1b). Bioinformatic analyses suggest that it influences the binding of transcription factors within a cis-regulatory element [6,7]. Both *HDAC9* and *TWIST1* have good evidence for being responsible for its effect on cardiovascular disease. *HDAC9* encodes Histone Deacetylase 9, which acts as a transcriptional and immune signalling regulator. Various lines of evidence, including mouse knockouts, suggest this has a role in atherosclerotic plaque growth and instability [8]. In cells related to immune function (peripheral blood mononuclear cells, primary macrophages and Jurkat cells), the rs2107595 risk allele associates with increased *HDAC9* but not *TWIST1* expression [6,9]. On the other hand, Twist-related protein 1, the protein encoded by *TWIST1*, is a transcription factor belonging to the basic helix-loop-helix family, which has been implicated in many biological pathways including mesodermal differentiation in embryonic development, the epithelial–mesenchymal transition process, tumour progression, fibrosis and skeletal development [10–15]. Single-cell ATAC-seq data indicate that, within vascular smooth muscle cells, rs2107595 physically interacts with the *TWIST1* promoter to increase *TWIST1* expression but not *HDAC9* expression [16], with implications for cell calcification and atherosclerotic plaque growth [7,17]. *TWIST1* may also have alternative effects in endothelial cells [18,19].

**Figure 1.**
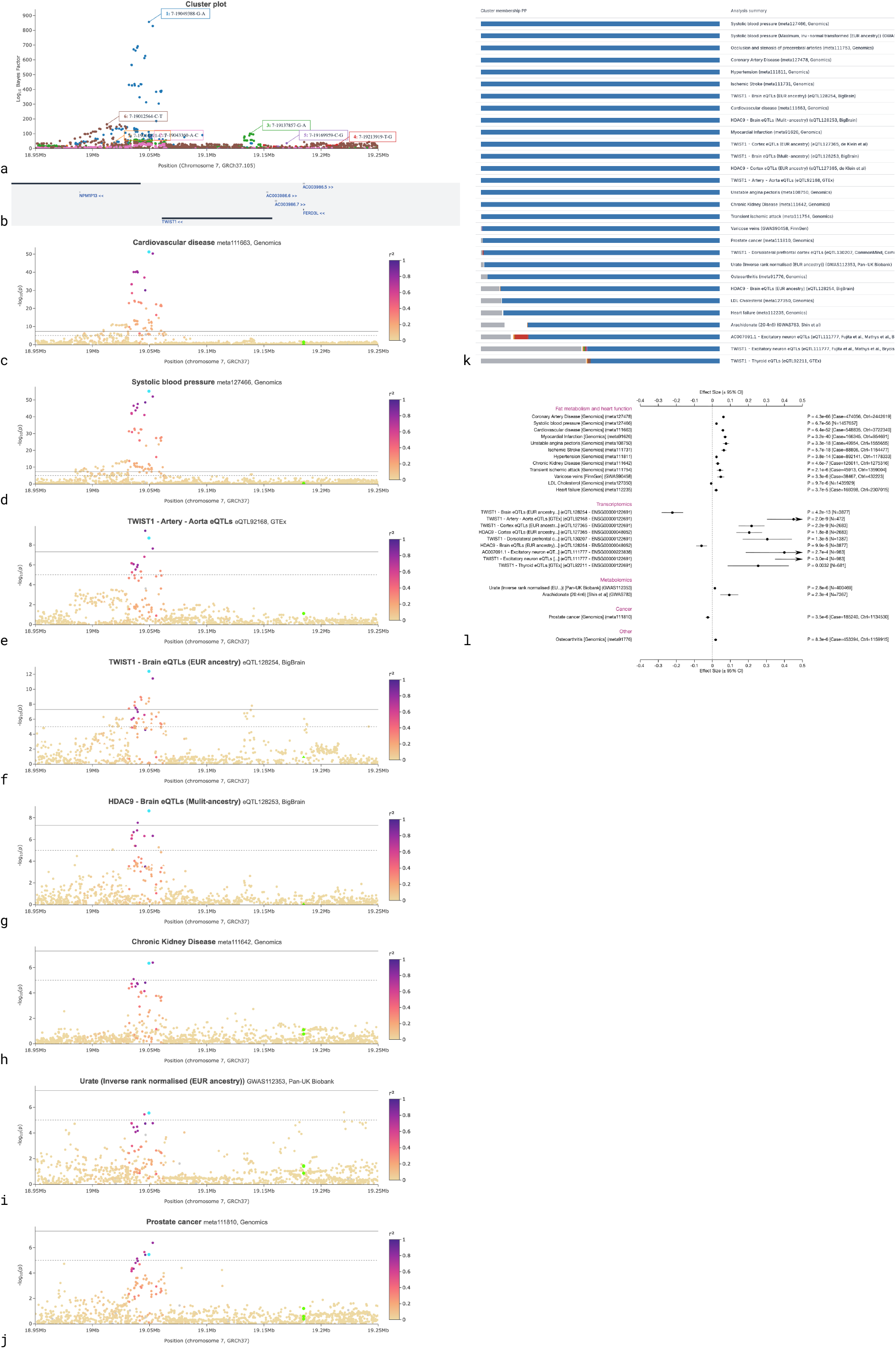
Multi-trait colocalisation analysis of the *HDAC9*-*TWIST1* locus using *MystraColoc*. Seven separate genetic signal clusters are inferred (**a**), for a 300kb region (Ensembl GRCh37.105) around *HDAC9*-*TWIST1* (**b**). Cluster 1 has 29 traits with a >0.5 posterior probability of cluster membership (**k**). Underlying association patterns (−log_10_p values) are shown for key traits of interest, with variants coloured by their LD (European ancestry 1000 Genomes resource) with the inferred lead variant (**c-j**); the blue dot indicates the Cluster 1 lead variant, rs2107595 located at chr7:19049388 (b37); solid and dotted horizontal lines indicate genome-wide (p<5×10^−8^) and suggestive (p<10^−5^) significance respectively. A forest plot of association effect sizes is shown for the lead variant rs2107595 (**l**), aligned to increasing allele count of the Alt allele.

We used *MystraColoc* to interrogate the extensive collection of GWAS analyses available in the Mystra platform. We input 411 datasets into *MystraColoc*, including 68 meta-analyses representing the largest available meta-analyses for all relevant major diseases, 83 other GWAS representing various molecular traits, 44 regional eQTL and 216 pQTL datasets. All datasets were required to have at least a moderately suggestive association signal, p<10^−6^, in a 300kb region around rs2107595. *MystraColoc* found 7 signal ‘clusters’ (Figure 1a). Cluster 1 was clearly identified as the cardiovascular trait cluster - 29 traits colocalised with a posterior probability >0.5 (Figure 1k), including many disease traits related to cardiovascular disease and its subtypes (coronary artery disease, hypertension, ischaemic stroke, myocardial infarction, and others). The underlying association signal for cardiovascular disease is shown in Figure 1c, indicating a very strong genome-wide significant signal (p-value ~10^−50^ for the cluster’s lead variant shown in blue). There is also a strong genome-wide significant signal for systolic blood pressure (Figure 1d), as one might expect given the presence of clinical hypertension as a trait in this cluster. Seven eQTLs for *TWIST1* are found by *MystraColoc* to colocalise with the cluster, and two for *HDAC9*. The *TWIST1* eQTLs implicate the artery and the brain (Figure 1e-f), whereas the *HDAC9* eQTL signals implicate only the brain (Figure 1g). Given the clear mechanistic hypothesis supported by previous work for arterial tissue to be the site of action for an influence on cardiovascular traits [16], our data support *TWIST1* as being the causal gene underpinning the associations for Cluster 1.

Thanks to the ability of *MystraColoc* to analyse weaker signals simultaneously, some interesting sub-genome-wide significant association signals for other traits also colocalise with Cluster 1 (Figure 1h-j). The direction of effect of the rs2107595 CVD-risk allele on these traits is shown in Figure 1l. The CVD risk allele associates with increased urate levels, chronic kidney disease and osteoarthritis risk, but decreased LDL cholesterol levels and prostate cancer risk. While both *HDAC9* and *TWIST1* may causally influence CVD risk via different pathways, it is notable that *TWIST1* is implicated in kidney fibrosis, skeletal pathology and cancer progression [10,12,14].

### *MystraColoc* colocalises more accurately

To quantify the performance of different muti-trait colocalisation approaches, we simulated repeated collections of 220 GWAS datasets in a single genomic region, designed to represent a range of effect sizes and sample sizes reflective of a mix of high-level traits, eQTLs and pQTLs. Three causal variants in mutually low LD were simulated for each collection. 201 datasets were simulated under an assumption of causality (100, 100 and 1 separate datasets for each causal variant, with different ancestries, effect sizes and sample sizes), and 19 GWAS datasets were simulated assuming no causal variant. See Methods for details.

For each collection of 220 GWAS datasets, we defined the sensitivity or true positive rate (TPR) as TP / (TP + FN), the false positive rate (FPR) as FP / (FP + TN), and the accuracy as (TP + TN) / (TP + FP + TN + FN), where TP is the number of dataset pairs that both truly shared a causal variant and were found to colocalise, FP is the number of pairs that did not truly share a causal variant and were found to colocalise, TN is the number of pairs that did not truly share a causal variant and were found not to colocalise, and FN is the number of pairs that did not truly share a causal variant and were found to colocalise.

Across the simulated runs, *MystraColoc* is more accurate than *HyPrColoc* (93.7% compared to 88.9%, a 5% increase overall, see Figure 2). The false positive rate is negligible for both methods (median 0% to 2 decimal places), so the differences in accuracy are accounted for by differences in the true positive rate (85.5% compared to 73.7%, an 8% increase overall).

**Figure 2.**
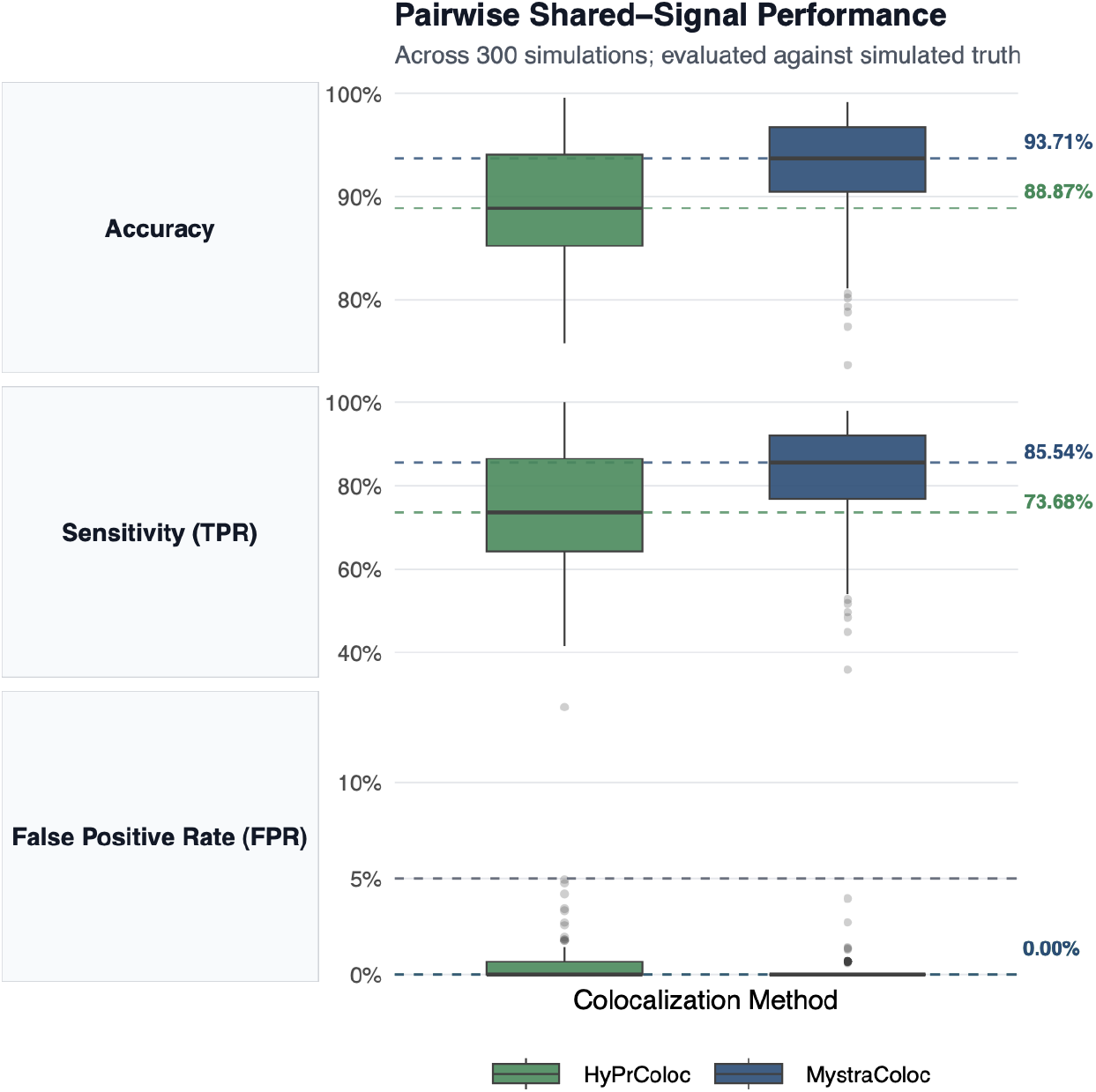
Accuracy, true and false positive rates of *HyPrColoc* and *MystraColoc* to identify dataset pairs with or without truly shared causal variants. Dashed lines indicate median values across simulations for *HyPrColoc* (green) and *MystraColoc* (blue).

### *MystraColoc* maintains performance even when the causal variant is in a high LD block

We separated results according to the number of LD tags (few [⩽5] / average [>5 and ⩽30] / many [>30] number of variants in high LD with the simulated causal variant, r^2^>0.9 in European ancestry 1000 Genomes resource). We found that *MystraColoc* maintains its accuracy across all LD patterns. In contrast, *HyPrColoc* performance is lower, with accuracy further reduced as the number of LD tags increases (Figure 3).

**Figure 3.**
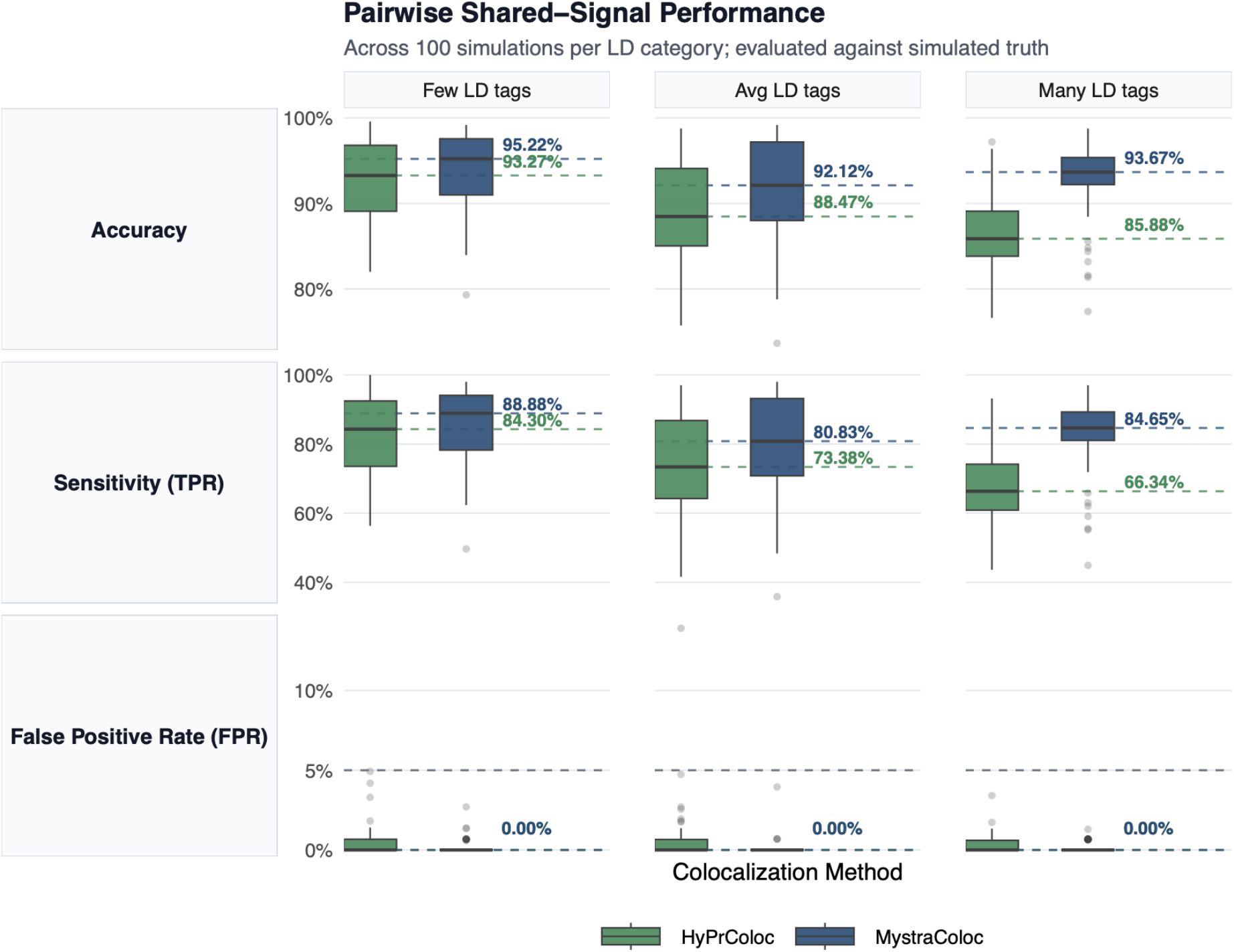
Accuracy, true and false positive rates of *HyPrColoc* and *MystraColoc* to identify dataset pairs with or without truly shared causal variants, with signals stratified by the number of LD tags (few [⩽5] / average [>5 and ⩽30] / many [>30] with r^2^>0.9 with the causal variant in European ancestry 1000 Genomes resource). Dashed lines indicate median values across simulations for *HyPrColoc* (green) and *MystraColoc* (blue).

### *MystraColoc* better identifies true cluster size and number

Focussing on the two causal variants with multiple datasets in our simulations, *MystraColoc* outperforms *HyPrColoc* by better recovering the true underlying cluster structure (Figure 4). Compared to the true number of 100 colocalising datasets, *MystraColoc* identifies a median of 88 datasets per cluster, whereas *HyPrColoc* only identifies a median of 39. Alongside this, *MystraColoc* correctly identifies the true number of clusters (two) in all the simulation runs. Thus, while *MystraColoc* is sometimes unable to assign weaker signals to a cluster, resulting in a median number slightly below the target of 100, *MystraColoc* properly captures the genetic architecture of the simulated data. In contrast, *HyPrColoc* generates an excess of clusters (median 5 clusters). This indicates a tendency of *HyPrColoc* to oversplit datasets into multiple clusters, obscuring the true colocalisation patterns.

**Figure 4.**
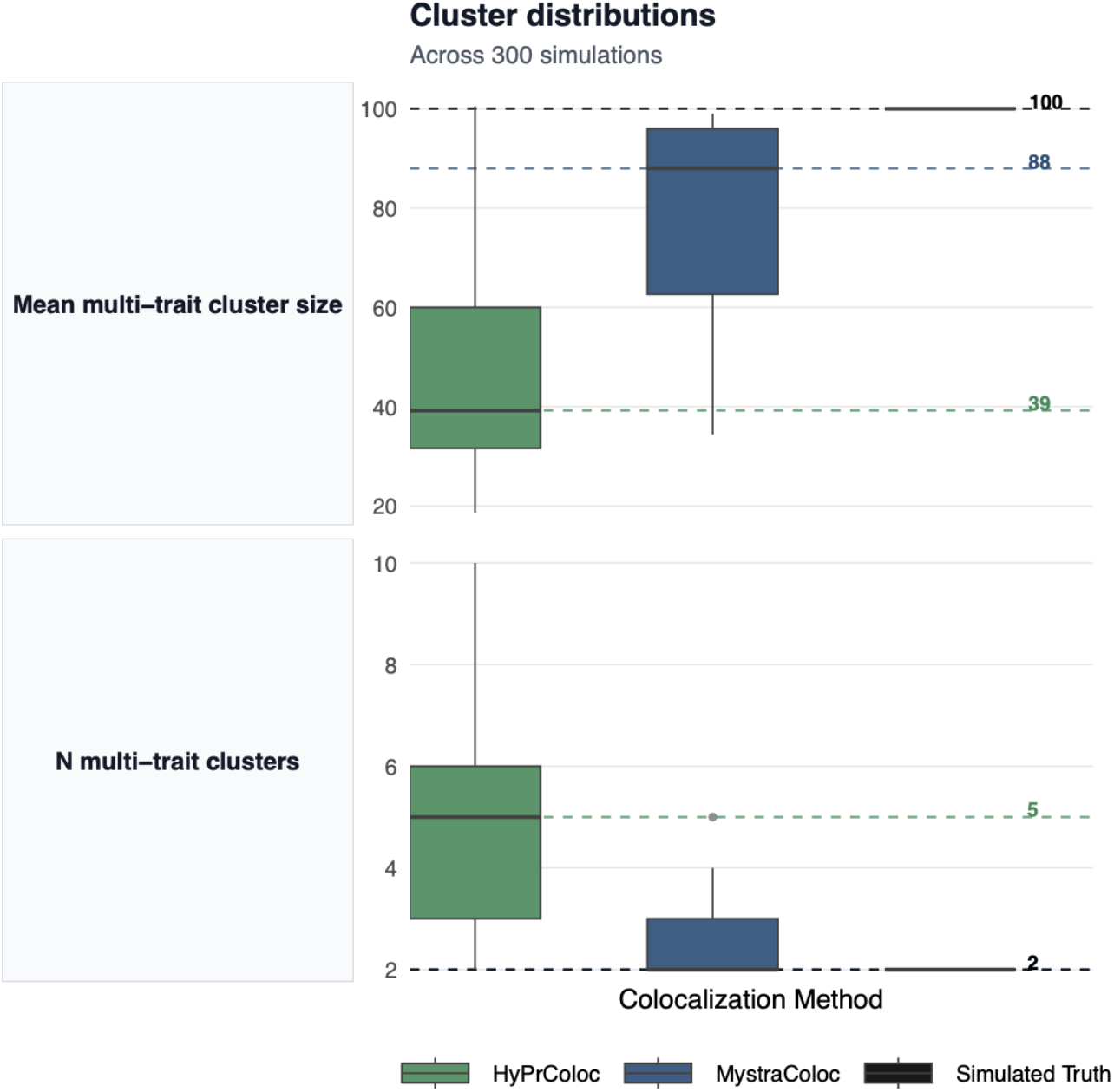
Distributions of number of multi-trait clusters and number of datasets per cluster for *HyPrColoc* and *MystraColoc*, compared to simulated truth (two simulated clusters of 100 studies each). Dashed lines indicate median values across simulations for *HyPrColoc* (green) and *MystraColoc* (blue) and simulated truth (black).

## Discussion

*MystraColoc* is an improved method for colocalising hundreds to thousands of traits. Multi-trait colocalisation unlocks the potential of large-scale GWAS platforms like Mystra, by revealing the shared biology implied by shared genetic signals. Taking one example, we show that the strong genome-wide significant signal for traits related to cardiovascular disease at the *HDAC9*-*TWIST1* locus can be linked to differential expression of *TWIST1* in arterial tissue, alongside other sub-genome-wide significant signals for urate, chronic kidney disease, osteoarthritis and prostate cancer.

Another widely used method for multi-trait colocalisation is *HyPrColoc*. We show via realistic simulations that *MystraColoc* outperforms *HyPrColoc*, both in its ability to correctly colocalise pairs of traits, even when there are multiple high LD tags for the true causal variant, and in its ability to recover the true underlying number of clusters and number of colocalising traits per cluster. *MystraColoc* achieves this via an efficient formulation of the relevant likelihoods, which allows the solution space to be explored more widely and effectively than the branch and bound approach used by *HyPrColoc*. Another advantage of *MystraColoc* is that it simultaneously colocalises shared traits and finemaps the causal variant, allowing for more accurate variant identification.

A remaining issue not tackled by either of the two algorithms presented here is the presence of multiple causal variants within a dataset, which should lead to a single dataset being assigned to multiple clusters. Neither *MystraColoc* nor *HyPrColoc* implement this option. A potential solution would be to use conditional analysis prior to multi-trait colocalisation to feed these signals as multiple datasets into the colocalisation algorithm. The ability of *MystraColoc* to scale to a large number of GWAS datasets makes it possible to chain these analytical tools at scale, and we are currently investigating these refinements.

In summary, *MystraColoc* unlocks the power of multi-trait colocalisation, and multi-trait colocalisation unlocks the power of Mystra.

## Methods

### MystraColoc

*MystraColoc* is an iterative Bayesian algorithm that simultaneously performs multi-trait colocalisation and finemaps the likely causal variants within multi-trait signal clusters. Within a given region of the genome, the model assumes that each GWAS dataset has either 0 or 1 causal variants. We refer to a group of datasets which are assumed to have the same causal variant as a cluster. The posterior distributions of both cluster membership (colocalisation) and casual variant location (finemapping) are jointly estimated by the algorithm. The algorithm uses Gibbs sampling to iteratively: 1) calculate the posterior distribution on the position of the causal variant for each cluster; 2) update the assignment of each dataset to either the null set (datasets with no causal variant), one of the existing clusters, or to a new cluster. The algorithm is implemented so that the prior distribution on the number and sizes of the clusters follows that of a Dirichlet random measure (also known as a Chinese Restaurant Process), with concentration parameter 0.01 [20]. The cluster likelihood follows the form described in [21], assuming that all trait effect sizes are independent both in terms of sampling variance and true effect variance. The prior probability of there being a causal variant for a given dataset in the region is set to 0.1, in line with the established view that most complex traits are highly polygenic. The prior probability of each variant being a causal variant for a given cluster is uniform on the number of variants in the region. The algorithm is run for sufficient iterations to allow convergence. The output of the algorithm is a posterior probability of cluster membership for each dataset (colocalisation), and the probability that each variant is causal in each of the clusters (fine-mapping).

### HyPrColoc

*HyPrColoc* (Hypothesis Prioritisation for multi-trait Colocalization) is a deterministic Bayesian method designed to identify shared genetic association signals from GWAS data across multiple traits [4]. *HyPrColoc* starts by approximating the posterior probability of colocalization through the enumeration of a small number of most likely potential causal configurations. Under the assumption that any single dataset possesses at most one causal variant in the region of interest, the algorithm calculates the Regional Association Probability (P_R_), which describes whether all traits share an association with one or more variants within the genomic region, and the Alignment Probability (P_A_), which describes whether these shared associations are due to a single shared causal variant, and multiplies these two together to arrive at the Posterior Probability of Full Colocalization (PPFC). If P_R_ is high but P_A_ is low, a branch and bound divisive clustering algorithm is used to split the GWAS set into subsets until a sufficiently large PPFC is achieved.

### GWAS simulation scheme

Regional GWAS datasets (sampled per-allele effect sizes and their standard errors) were simulated based on the linkage disequilibrium (LD) and minor allele frequency (MAF) distributions of a real region of the human genome (500kb region around the gene *CTLA4* on chromosome 2; LD and MAF obtained from the 1000 Genomes resource [22]), using a simulation scheme based on the same methodology as GWASBrewer [23]. Variants with MAF>1% in European ancestry were selected for inclusion in the GWAS datasets. Three variants (V1, V2 and V3) were selected at random to be causal variants, conditional on low LD between these variants (r^2^<0.1 in European ancestry). 100 GWAS datasets were simulated with V1 as their causal variant; a separate 100 GWAS datasets were simulated with V2 as their causal variant; a separate single GWAS was simulated with V3 as its causal variant; and finally 19 GWAS datasets were simulated without any causal variant. Study sample sizes chosen to reflect a variety of realistic study types: 100k for high-level traits (HLT); 50k for pQTLs; and 100 for eQTLs (typical of some GTEx tissues). All traits were assumed to be quantitative, but it can be noted that a 50:50 case-control GWAS with a four times greater total sample size would generate equivalent effect size standard errors [24].

Effect sizes were randomly selected from normal distributions to reflect one of three significance types: suggestive (sug, 10^−5^>p>10^−7^); genome-wide significant (gws, 10^−7^>p>10^−20^); and extreme (ext, 10^−200^<p<10^−300^). V1 and V2 causal datasets were generated as follows: 15 HLT_sug; 20 HLT_gws; 5 HLT_ext; 10 pQTL_sug; 10 pQTL_gws; 10 pQTL_ext; 15 eQTL_sug; and 15 eQTL_gws. The V3 causal dataset was generated as an HLT_gws.

100 separate GWAS collections were generated, each consisting of 220 GWAS datasets. The majority of datasets were simulated using European ancestry LD and MAF distributions, but 15 datasets within the V1 and V2 causal datasets respectively (30 in total) were simulated using Sub-Saharan African ancestry LD and MAF distributions.

## Acknowledgments

The implementation of *MystraColoc* in the Mystra platform is the result of a large collaborative effort across the Science, Data and Software Engineering, Product and other teams at Genomics Ltd. The *MystraColoc* algorithm was originally developed by Chris C. A. Spencer.

